# S-Palmitoylation-Dependent Regulation of Cardiomyocyte Rac1 Signaling Activity and Cardiac Hypertrophy

**DOI:** 10.1101/2021.05.14.444015

**Authors:** Matthew J. Brody, Tanya A. Baldwin, Arasakumar Subramani, Onur Kanisicak, Ronald J. Vagnozzi, Jeffery D. Molkentin

**Author notes:** corresponding authors Correspondence: Jeffery D. Molkentin, Matthew J. Brody.

## Abstract

S-palmitoylation is a reversible lipid modification that regulates trafficking, localization, activity, and/or stability of protein substrates by serving as a fatty acid anchor to cell membranes. However, S-palmitoylation-dependent control of signal transduction in cardiomyocytes and its effects on cardiac physiology are not well understood. We performed an in vivo gain-of-function screen of zinc finger Asp-His-His-Cys (zDHHC) family S-acyl transferases that catalyze S-palmitoylation and identified the Golgi-localized enzyme zDHHC3 as a critical regulator of cardiac maladaptation. The closely-related enzyme, zDHHC7, also induced severe cardiomyopathy but this effect was not observed with overexpression of plasma membrane enzyme zDHHC5, endoplasmic reticulum enzyme zDHHC6, or Golgi enzyme zDHHC13. To identify effectors that may underlie zDHHC3-induced cardiomyopathy we performed quantitative site-specific S-acyl proteomics in zDHHC3-overexpressing cells that revealed the small GTPase Rac1 as a novel substrate. We generated cardiomyocyte-specific transgenic mice overexpressing zDHHC3, which develop severe cardiac disease. Cardiomyopathy and congestive heart failure in zDHHC3 transgenic mice are preceded by enhanced S-palmitoylation of Rac1 and induction of additional Rho family small GTPases including RhoA, Cdc42, and the Rho family-specific chaperone RhoGDI. In contrast, transgenic mice overexpressing an enzymatically-dead mutant of zDHHC3 do not exhibit this profound induction of RhoGTPase signaling or develop cardiac disease. Rac1 S-palmitoylation, plasma membrane localization, activity, and downstream hypertrophic signaling were substantially increased in zDHHC3 overexpressing hearts. Taken together, these data suggest inhibition of zDHHC3/7 S-acyl transferase activity at the cardiomyocyte Golgi or disruption of Rac1 S-palmitoylation as novel therapeutic strategies to treat cardiac disease or other diseases associated with enhanced RhoGTPase signaling.

## INTRODUCTION

Diverse intracellular signaling mechanisms participate in cardiac homeostasis and physiologic growth as well as maladaptation in cardiac pathologic hypertrophy and heart failure ^1-3^. Pathological signaling in cardiomyocytes is often transduced from the sarcolemma (plasma membrane) by GTPases that activate downstream intracellular signaling cascades ^4-10^. Activation of small GTPases is dynamically regulated by guanine nucleotide dissociation inhibitors (GDIs), GTPase activating proteins (GAPs), and guanine nucleotide exchange factors (GEFs) ^11-14^. Additionally, some small GTPases such as H-Ras, N-Ras, and Ras-related C3 botulinum toxin substrate 1 (Rac1) undergo S-palmitoylation or S-acylation, a reversible lipid modification on cysteine residues that governs their dynamic association with the plasma membrane and subsequent activation of downstream effectors ^15-17^. Moreover, certain GTPase regulatory proteins, including p63 RhoGEF ^18^ and the regulator of G-protein signaling (RGS) proteins that function as GAPs for heterotrimeric Gα subunits ^19, 20^ are S-palmitoylated, providing another layer of S-palmitoylation-dependent control of signaling by G-proteins. However, the enzymes controlling fatty acylation of GTPases and their regulators, and consequences of S-palmitoylation on signaling by small GTPases in vivo, are not well-established, particularly in the context of cardiomyocyte signaling in hypertrophy and heart failure.

Cardiomyocyte-specific overexpression of RhoA or Rac1 causes cardiac failure in mice ^9, 21^ and RhoGTPase signaling is activated in murine cardiomyopathy ^6, 22^ and human heart failure ^23^. Rac1 in particular is an essential mediator of reactive oxygen species (ROS) generation in the heart through its role as a regulatory subunit of the nicotinamide adenine dinucleotide phosphate (NADPH) oxidase-2 (Nox2) complex ^24, 25^ and is required for cardiac hypertrophy and oxidative stress in response to angiotensin II ^5^. Importantly, impairment of Rac1 activity and oxidative stress are primary mechanisms of statin-mediated cardioprotection in animal models ^5, 26, 27^ and statin treatment ameliorates Rac1 activation, NADPH oxidase activity, and ROS production in the failing human heart ^23^. However, the mechanisms that modulate Rac1 signaling in the heart remain ill-defined.

Dysregulation of S-palmitoylation has recently emerged as an important molecular participant in human disease ^28, 29^. The addition of a palmitoyl moiety at cysteine residues is reversible and sets it apart from other post-translational lipid modifications ^28, 30^. The dynamic nature of S-palmitoylation provides a powerful regulatory mechanism akin to protein phosphorylation, with diverse effects on protein localization and function. Small GTPases and G-protein α subunits undergo rapid cycles of S-palmitoylation and depalmitoylation to elicit sustained signaling activity ^31, 32^, implicating S-palmitoylation as a nodal regulatory point controlling intracellular signal transduction. There are 23 zinc finger Asp-His-His-Cys (zDHHC) S-acyltransferases (encoded by the *Zdhhc* genes) in mammals that catalyze S-palmitoylation ^33-35^, which is reversed predominantly by two major cytosolic depalmitoylases, acyl protein thioesterase 1 and 2 (APT-1 and APT-2 encoded by the *Lypla1* and *Lypla2* genes) ^30^ along with three serine hydrolases of the α/β hydrolase domain-containing protein 17 (ABHD17) family that also exhibit depalmitoylase activity ^36, 37^. zDHHC S-acyltransferases are polytopic transmembrane proteins, many of which localize to the endoplasmic reticulum or Golgi apparatus in addition to the plasma membrane, endomembrane system, or intracellular vesicles ^33, 38^. Despite some common substrates amongst different zDHHC enzymes, there is generally strong selectivity and substrate specificity imparted by substrate recruitment domains on the cytoplasmic tails of zDHHCs, zDHHC interaction domains on protein substrates, and restricted subcellular and tissue and cell-type expression patterns of zDHHC enzymes and their substrates ^30, 39, 40^. Even amongst Golgi-localized zDHHCs there is specified recruitment of substrates by their cognate zDHHC enzyme ^41, 42^. Thus, S-palmitoylation is a tightly controlled regulatory mechanism that underlies intracellular signal transduction via dynamic targeting of proteins to membrane microdomains.

Modern techniques to detect S-palmitoylated proteins ^31, 43, 44^ and the discovery of zDHHC enzymes that catalyze this form of lipidation ^39, 41^ have greatly accelerated progress in understanding physiologic functions of S-palmitoylation and led to the recent identification of zDHHC-regulated mechanisms in disease. For example, the Golgi enzyme zDHHC13 palmitoylates melanocortin-1 receptor (MC1R) to promote melanin production and protect from UV-induced melanoma ^45, 46^ while the plasma membrane enzyme zDHHC20 modifies the epidermal growth factor receptor (EGFR) ^47-49^. Loss of EGFR S-palmitoylation or zDHHC20 sensitizes cancer cells to EGFR inhibitors ^48^ and impairs phosphoinositide 3-kinase (PI3K) signaling in K-Ras mutated cancers ^49^. Moreover, S-palmitoylation of signal transducer of activator of transcription 3 (STAT3) ^50^ and stimulator of interferon genes (STING) ^51-53^ are fundamental signaling mechanisms in the pathogenesis of inflammatory diseases that can be effectively targeted in pre-clinical models for the treatment of colitis and Aicardi-Goutiéres syndrome, respectively. Collectively, these studies have uncovered S-palmitoylation-dependent signal transduction mechanisms that are being pursued as novel therapeutic drug targets ^45, 52, 53^. However, studies of zDHHC enzymes and S-palmitoylation in the heart are largely limited to investigation of ion channel regulation and electrophysiology ^40, 54^ and roles of S-palmitoylation in cardiomyocyte signal transduction and hypertrophy and heart failure have not been fully evaluated.

Here, we performed an in vivo screen using recombinant adeno-associated virus (AAV)-mediated overexpression, which identified the Golgi-localized enzyme zDHHC3 as a potent inducer of cardiac maladaptation, decompensation, and heart failure. While zDHHC3 is expressed in the heart ^38, 40^, much of the prior work in the field focused on its function in neurons ^55-58^ and its functions in the cardiomyocytes are not known. We observed that Rac1 is a novel substrate of zDHHC3 using an unbiased proteomic approach and that cardiomyocyte-specific transgenic mice overexpressing zDHHC3 develop profound cardiomyopathy and heart failure. This disease phenotype is preceded by enhanced Rac1 S-palmitoylation and associated with dramatically enhanced plasma membrane localization, protein levels, and activity of Rac1 and other Rho family GTPases. These studies identify zDHHC3 S-acyltransferase activity at the cardiomyocyte Golgi as a nodal regulator of RhoGTPase activity and cardiomyopathy in association with regulated S-palmitoylation of Rac1.

## METHODS

### Animals

Cardiomyocyte-specific transgenic mice overexpressing zDHHC3 were generated by subcloning mouse *Zdhhc3* cDNA (Dharmacon, #MMM1013-202763213) into the re-engineered tetracycline inducible α-myosin heavy chain (α-MHC) promoter expression vector that permits tetracycline/doxycycline extinguishable expression in the presence of a second transgene expressing the tet-transactivator (tTA) expressed by the unmodified α-MHC promoter expression vector ^59^. The DNA construct was digested with Not I restriction endonuclease and the promoter-cDNA fragment gel purified for oocyte injection at the Cincinnati Children’s Hospital Transgenic Animal and Genome Editing Core Facility as described previously ^60, 61^. Enzymatically-dead zDHHC3 mutant transgenic mice overexpressing zDHHC3^DHHS^ in cardiomyocytes were generated by site-directed mutagenesis of the α-MHC-*Zdhhc3* promoter-transgene construct using the QuikChange II XL Site-Directed Mutagenesis Kit (Agilent) to encode a mutation of Cys-157 in mouse zDHHC3 protein to Ser. Primers used for mutagenesis were Forward 5’-GCAAGATGGATCACCACAGTCCTTGGGTCAACAAC-3’ and Reverse 5’-GTTGTTGACCCAAGGACTGTGGTGATCCATCTTGC-3’. Transgenic mice were generated on the FVB/N genetic background. To induce transgene expression in the adult heart, transgenic mice were bred on doxycycline-containing chow (625 mg/kg diet, Cincinnati Lab Supply, #TD1811541) to repress transgene expression until 3 weeks of age when experimental mice were weaned from the dams and placed on a normal lab chow diet. All molecular analyses were performed in the high-expressing line of zDHHC3 transgenic mice two months following induction of transgene expression (removal of doxycycline) in the adult heart unless otherwise stated.

Adeno-associated virus serotype 9 (AAV9) was generated by subcloning full-length mouse *Zdhhc* cDNAs (kind gift of Dr. Masaki Fukata, National Institute for Physiological Sciences, Japan) ^41^ with 2x hemagglutinin (HA) epitopes on the N-terminus into the pAAV-MCS vector (Agilent) and AAV9 was produced by Vigene or the Howard Hughes Medical Institute Viral Vector Core. Mouse pups were injected in the chest cavity at postnatal day 6 with 1 × 10^12^ viral genomes of the indicated AAV9 as described previously ^62^ with the exception of Zdhhc3 and Zdhhc7 that were injected at lower doses of 0.5 × 10^12^ or 1 × 10^11^ viral genomes per pup, respectively, due to lethality associated with robust expression. Controls were injected with 1 × 10^12^ viral genomes of empty AAV9 vector or sterile PBS. AAV studies were performed in CD1 mice with the exception of Zdhhc7 and associated controls in supplemental Fig 1 that were performed in the FVB/N strain.

Echocardiography was performed as described previously ^63, 64^. Noninvasive electrocardiography was performed on conscious mice using the ECGenie Awake Lab Animal ECG System (Mouse Specifics, Inc.) and analyzed with LabChart software (ADI Instruments). Mortality was defined as a mouse being found dead in the cage or veterinarian-recommended euthanasia due to symptoms of congestive heart failure. All animal procedures were approved by the Cincinnati Children’s Hospital Institutional Animal Care and Use Committee (IACUC) and conformed to the Guide for the Care and Use of Laboratory Animals (NIH).

### Western Blotting, Immunoprecipitations, Membrane Fractionation, and GTPase Activity

For evaluation of small GTPase activity and small GTPase protein levels, hearts were homogenized in assay buffer (25 mM HEPES pH 7.5, 150 mM NaCl, 1% NP-40, 10 mM MgCl_2_, 1 mM EDTA, and 2% glycerol) with protease inhibitors (Roche) and lysates were cleared by centrifugation. RhoA activity was evaluated by affinity purification of RhoA-GTP using rhotekin agarose beads (Cell Biolabs) and the activity of Rac1 and Cdc42 were assessed by affinity purification using magnetic beads coupled to the p21-binding domain of p21-activated protein kinase (PAK) (Millipore) that specifically binds the active (GTP-bound) forms of Rac1 and Cdc42. Following affinity purification, GTP-bound small GTPases were eluted from beads for SDS-PAGE by boiling in Laemmli buffer.

Western blotting was performed as described previously ^63, 65^. Mouse hearts were homogenized in radioimmunoprecipitation assay (RIPA) buffer (50 mM Tris•HCl pH 7.4, 1% triton X-100, 1% sodium deoxycholate, 1 mM EDTA, 0.1% SDS) or, for detection of zDHHC proteins, in 50 mM Tris•HCl pH 7.6, 10 mM Na_4_PO_2_O_7_•10H_2_O, 6 M urea, 10% glycerol, and 2% SDS containing Halt protease and phosphatase inhibitor cocktail (Thermo Scientific) and then sonicated, clarified by centrifugation, and boiled in Laemmli buffer. Biochemical fractionation of mouse hearts into membrane and cytosolic fractions was performed exactly as described elsewhere ^66^. HA-tagged zDHHC proteins were immunoprecipitated from cardiac lysates with anti-HA magnetic beads (Pierce, #88836) and eluted by boiling in Laemlli buffer. Samples were separated by SDS-PAGE and transferred to polyvinylidene difluoride (PVDF) membranes (Millipore Immobilon-FL, #IPVH00010) for immunoblotting. PVDF membranes were blocked in 5% dry milk diluted in Tris-buffered saline with 0.1% tween-20 (TBST), incubated with primary antibodies diluted in 5% milk in TBST overnight at 4°C followed by incubation with LiCor IRDye secondary antibodies diluted 1:10,000 in 5% milk in TBST with 0.02% SDS for two hours at room temperature, and imaged on a Li-Cor Odyssey CLx imaging system. Primary antibodies used were RhoA (Cell Signaling Technology, #2117, 1:500), Rac1 (BD Transduction Laboratories, #610650, 1:500), Cdc42 (Abcam, #ab64533, 1:500), RhoGDIα (BD Transduction Laboratories, #610255, 1:4,000), zDHHC3 (Abcam, #ab31837, 1:500), zDHHC7 (Abcam, #ab138210, 1:500), p-extracellular signal-regulated kinase (ERK) 1/2 (Cell Signaling Technology, #4370, 1:500), ERK1/2 (Cell Signaling Technology, #9102, 1:500), pan-Ras (Thermo Fisher, #MA1-012, 1:1,000), HA (Abcam, #ab9110, 1:1,000), PAK1 (Cell Signaling Technology, #2602, 1:500), H-Ras (Santa Cruz Biotechnology, #sc-29, 1:500), integrin β1D (Millipore, #MAB1900, 1:1,000), tubulin (Sigma, #T5168, 1:1,000), and Gapdh (Fitzgerald, #10R-G109A, 1:50,000).

### Immunocytochemistry

Immunocytochemistry was performed on adult cardiomyocytes in suspension exactly as described previously ^61, 64^. Cardiomyocytes were isolated from mouse hearts by Langendorff perfusion, fixed with 4% paraformaldehyde for 15 minutes at room temperature, incubated in blocking solution (PBS, 5% goat serum, 1% BSA, 1% glycine, 0.2% triton X-100) for one hour at room temperature and then immunostained with Rac1 (BD Transduction Laboratories, #610650) or zDHHC3 (Abcam, #ab31837) antibodies. Primary antibodies diluted 1:50 in blocking solution overnight at 4°C. Cardiomyocytes were then washed in PBS with 0.1% NP-40, incubated with Alexa Fluor secondary antibodies (Molecular Probes) diluted 1:1,000 in blocking solution for two hours at room temperature, washed again in PBS with 0.1% NP-40, and mounted on slides with Prolong Diamond Antifade Mountant with 4′,6-diamidino-2-phenylindole (DAPI) (Molecular Probes). Imaging was performed using a Nikon A1 Confocal microscope.

### Acyl Resin-Assisted Capture and Mass Spectrometry

S-palmitoylated proteins were purified from cardiac lysates for immunoblotting by Acyl Resin-Assisted Capture (Acyl-RAC) as described previously ^43^. Briefly, cardiac lysates were made in RIPA buffer as described above, diluted with 100 mM HEPES pH 7.4, 1 mM EDTA to a concentration of 2.5% SDS and free thiols were blocked with 0.2% methyl methanethiosulfonate (MMTS) at 42°C for 20 minutes. Proteins were then acetone precipitated at −20°C and samples centrifuged for 10 minutes at 10,000 x g. Protein pellets were washed four times in 70% ice cold acetone to remove excess MMTS and protein pellets were dried and solubilized in 100 mM HEPES pH 7.4, 1% SDS, and 1 mM EDTA with protease inhibitors at 37°C. Protein concentration was quantified and samples diluted to an equal concentration. For affinity purification of S-palmitoylated proteins 450 μL lysate was combined with 200 μL of 100 mM HEPES pH 7.4 with 1 mM EDTA, 300 μL of 1M NH_2_OH pH 7.4 or 150 mM Tris•HCl pH 7.4 as a negative control, and 30 μL thiopropyl sepharose (Sigma) and incubated at room temperature for three hours. Thiopropyl sepaharose beads were then washed four times in 100 mM HEPES pH 7.4, 0.3% SDS with 1 mM EDTA and S-palmitoylated proteins were eluted from thiopropyl sepharose by boiling in Laemmli buffer.

Stable 3T3 cells (ATCC) overexpressing zDHHC3 or green fluorescent protein (GFP) as a control were generated using the pLVX lentiviral system (Clontech). Cells were labeled by stable isotope labeling with amino acids in cell culture (SILAC) by passaging at least 9 times in Dulbecco’s Modified Eagle Medium (DMEM) for SILAC lacking lysine and arginine (Thermo Scientific) containing 10 % dialyzed fetal bovine serum (Thermo Scientific) and supplemented with [^13^C_6_, ^15^N_2_] L-lysine and [^13^C_6_, ^15^N_4_] L-arginine (Thermo Scientific) for “heavy” 3T3-zDHHC3 cells or normal L-lysine and L-arginine (Thermo Scientific) for “light” 3T3-GFP cells. Mass spectrometry sequencing of S-palmitoylated peptides was performed essentially as described previously ^43^ with slight modifications. Protein extracted from SILAC-labeled 3T3-Godz and 3T3-GFP cells was mixed 1:1 and S-palmitoylated proteins were purified by Acyl-RAC as described. Following the final wash, thiopropyl sepharose beads were incubated overnight at 37°C with 2 μg trypsin Gold (Promega, #V5280) in 50 mM NH_4_HCO_3_, 1mM EDTA. After on-resin trypsin digestion, thiopropyl sepharose beads were washed five times in 100 mM HEPES, 1% SDS, 1 mM EDTA then washed four times in 10 mM NH_4_HCO_3_ and once in 50 mM NH_4_HCO_3_. Captured peptides were then eluted from thiopropyl sepharose by incubation with 100 mM DTT (Roche) in 50 mM NH_4_HCO_3_ at 70°C for 45 minutes and further processed for mass spectrometry sequencing at the University of Cincinnati Proteomics Laboratory ^67^. Eluted peptides were alkylated with 200 uL of 400 mM iodoacetamide in 25 mM NH_4_HCO_3_ at 37°C for 2 hours and loaded onto C18 stage tips made from 3M Empore extraction disks, washed twice with 50 μl of 0.1% formic acid, and stage tips were then eluted three times with 50 μl of 80% acetonitrile/0.1% formic acid by centrifuging through the column at 1,600 x g for 5 min. Elutions were pooled, dried, and reconstituted in 6 ul of 0.1% formic acid and peptides were sequenced buy nanoscale liquid chromatography coupled to tandem mass spectrometry (nano LC-MS/MS) and searched using the Protein Pilot program (Sciex).

### Statistical Analyses

All statistical analyses were performed using GraphPad Prism with statistical testing as described in the figure legends.

## RESULTS

We performed an in vivo screen by overexpressing several DHHC enzymes in the heart with adeno-associated virus serotype 9 (AAV9). Pups were injected with AAV9 at postnatal day 6 to induce cardiac expression of the Golgi-localized enzymes zDHHC3 and zDHHC13 as well as the plasma membrane enzyme zDHHC5 and the endoplasmic reticulum-localized enzyme zDHHC6, and cardiac morphology and function were assessed one month later (Figs 1A, B). Enhanced expression of the Golgi-resident S-acyltransferase zDHHC3 specifically resulted in a profound cardiomyopathy that was not observed with overexpression of any of the other zDHHC enzymes tested (Fig 1C-E). Cardiac overexpression of zDHHC3 resulted in cardiac enlargement including ventricular and atrial dilation (Fig 1C), cardiac hypertrophy (Fig 1D), and substantial cardiac dysfunction indicative of cardiomyopathy (Fig 1E). The most homologous S-acyltransferase to zDHHC3 is another Golgi-localized enzyme, zDHHC7, and AAV9-mediated overexpression of this enzyme in the heart similarly induced severe cardiomyopathy within 3 weeks (Supplemental Fig 1). These data collectively suggest the activity of zDHHC3 and zDHHC7 at the cardiomyocyte Golgi promote pathogenic intracellular signaling that results in cardiac decompensation. Importantly, endogenous protein levels of both zDHHC3 and zDHHC7 were increased in the heart in response to pressure overload-induced hypertrophic stimulation (Fig 1F), suggesting a physiologic role of S-palmitoylation mediated by zDHHC3 and zDHHC7 in cardiac maladaptation.

**Figure 1.**
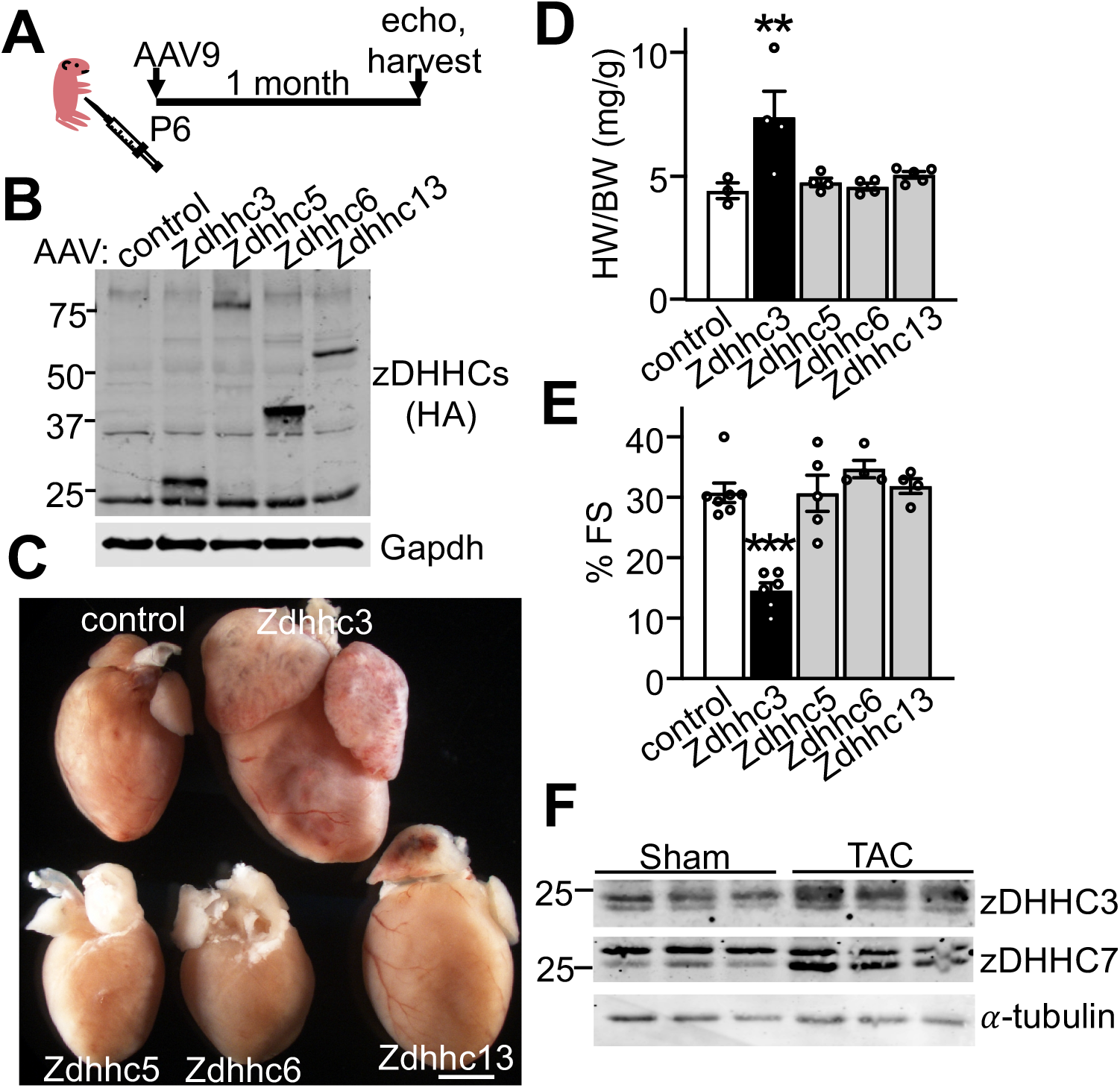
Enhanced activity of the Golgi-resident enzyme zDHHC3 causes cardiomyopathy. (A) Experimental design schematic and (B) Western blotting for AAV9-mediated overexpression of HA-tagged zDHHC enzymes in the heart. (C) Gross morphology, (D) heart weight-to-body weight ratios (HW/BW), n=3-5, (E) and fractional shortening (FS) as assessed by echocardiography in mice with cardiac overexpression of the indicated zDHHC enzymes. n=4-7. (F) Protein levels of zDHHC3 and zDHHC7 in mouse hearts after 8 weeks of pressure overload (TAC, transverse aortic constriction) compared to sham controls. Scale bar = 1 mm in C. **P<0.005, ***P<0.0001 compared to controls, ANOVA with Dunnett’s multiple comparisons test.

To further evaluate functions of zDHHC3-mediated S-palmitoylation in vivo, we generated mice with cardiomyocyte-specific overexpression of zDHHC3 using the binary regulated system consisting of the tetracycline transactivator (tTA) protein and the tet operator downstream of a modified α-myosin heavy chain (α-MHC) promoter ^59^ such that double transgenic mice (DTgZdhhc3) containing both the tTA and Zdhhc3 transgenes express zDHHC3 protein in the heart if doxycycline (Dox) is not present in the diet (“tet-off” system) ^68^ (Fig 2A, B). As an additional control we generated cardiac-specific transgenic mice that overexpress an enzymatically-dead zDHHC3 protein containing a Cys-to-Ser point mutation in its enzymatic DHHC domain (DTgZdhhc3^DHHS^) (Fig 2A, B). Western blotting of heart extracts from adult mice showed abundant overexpression of each protein in the heart compared with tTA controls (Fig 2B). Immunocytochemistry in isolated adult cardiomyocytes revealed the expected Golgi localization pattern for overexpressed wildtype and transferase-dead zDHHC3 proteins (Fig 2C). Transgenic mice on a normal diet with overexpression of zDHHC3 starting around birth (when ventricular α-MHC expression begins), but not mice overexpressing the enzymatically-dead zDHHC3^DHHS^ mutant, exhibited substantial mortality in young adulthood due to severe dilated cardiomyopathy with a median survival of 6 weeks of age in the low expressing lines (Fig 2D, E) and around 3 weeks of age in the high-expressing lines (data not shown). Gross morphological and histological analyses revealed dramatic cardiac enlargement and ventricular and atrial dilation in DTgZdhhc3 hearts (Fig 2E) and heart weight-to-body weight ratios confirmed significant cardiac hypertrophy (Fig 2F). Electrocardiography indicated severe bradycardia in DTgZdhhc3 mice (Supplemental Fig 2). Cardiac function and structure were evaluated in DTgZdhhc3 and DTgZdhhc3^DHHS^ mice by echocardiography and revealed significant left ventricular dilation (Fig 2G), systolic dysfunction (Fig 2H), and impaired cardiac contraction and cardiomyopathy (Fig 2I) in mice with cardiomyocyte-specific overexpression of zDHHC3 but not in mice expressing the zDHHC3^DHHS^ mutant (Figs 2G-I, Supplemental Table 1), demonstrating enhanced zDHHC3 S-acyltransferase activity in cardiomyocytes causes severe lethal dilated cardiomyopathy.

**Figure 2.**
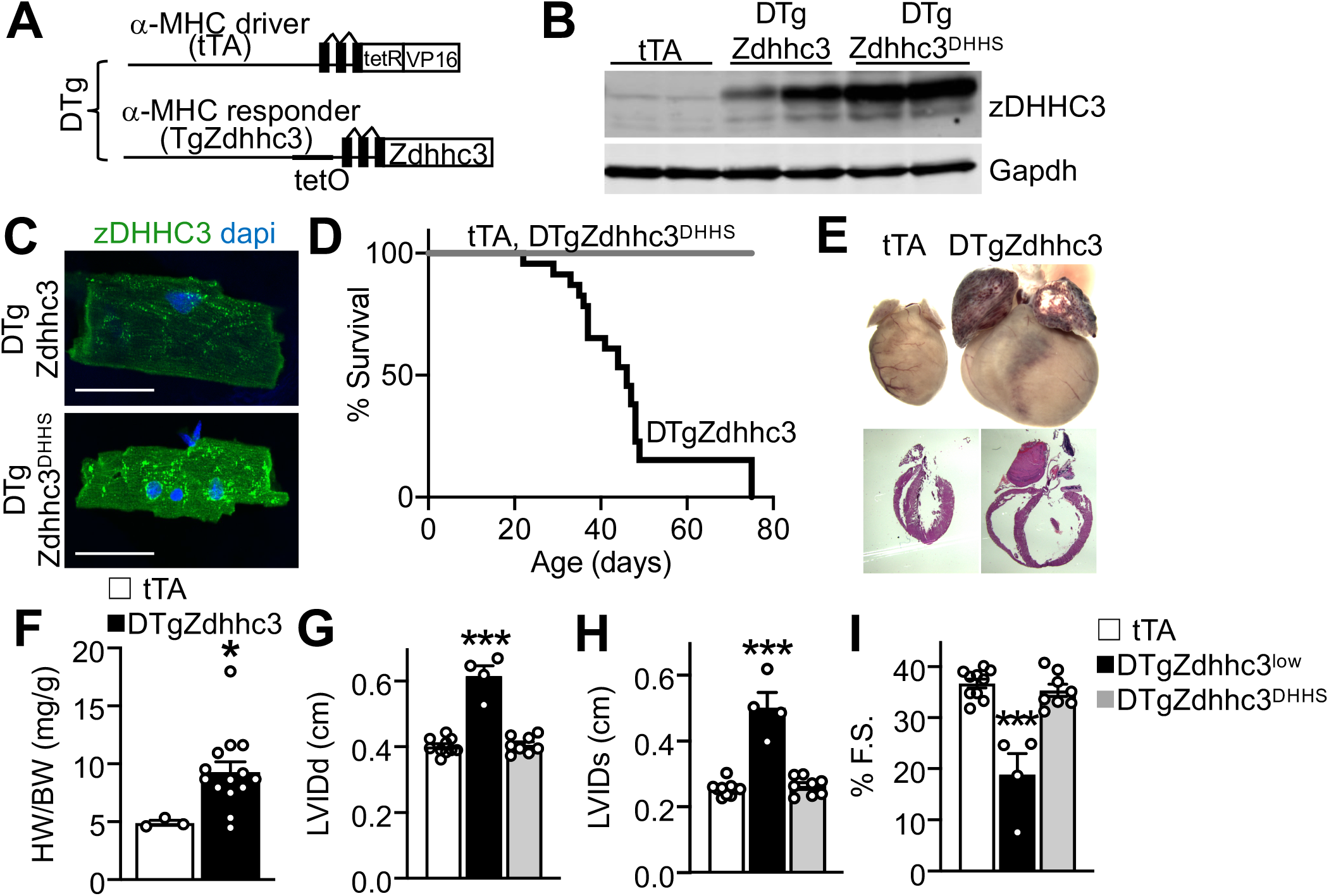
Cardiac-specific overexpression of zDHHC3 results in cardiomyopathy and lethality. (A) Schematic of binary “tet-off” inducible system used to overexpress zDHHC3 in the heart. (B-C) Cardiomyocyte-specific transgenic expression of zDHHC3 or zDHHC3^DHHS^ two months after doxycycline chow removal/transgene expression in the indicated high-expressing lines of mice. (B) Western blot for zDHHC3 expression in cardiac lysates. Gapdh is run as a loading and protein processing control. (C) Immunocytochemistry on adult isolated cardiomyocytes demonstrating Golgi localization pattern for both zDHHC3 and enzymatically-dead zDHHC3 ^DHHS^ protein in double transgenic (DTg) hearts. Scale bar = 50 μm. (D-I) Phenotypic characterization of low-expressing line of transgenic mice bred and maintained on normal lab chow to induce cardiomyocyte-specific expression of zDHHC3 from around birth. (D) Kaplan-Meier survival curve showing mortality in transgenic mice with perinatal cardiomyocyte-specific expression of zDHHC3. n=14 tTA, 21 DTgZdhhc3, and 13 DTgZdhhc3 ^DHHS.^ (E) Gross morphology and histology (H&E) of transgenic hearts at 7 weeks of age. (F) Heart weight-to-body-weight ratios at 4 weeks of age. n=3-14. *P<0.05 compared to tTA, unpaired t-test. (G-I) Echocardiographic measurement of (G) diastolic left ventricular inner diameter in diastole (LVIDd), (H) systolic LVID (LVIDs), and (I) % fractional shortening (FS) at 6-8 weeks of age. n=4-10. ***P<0.0001 compared to tTA, ANOVA with Dunnett’s multiple comparisons test.

To examine the role of zDHHC3 in the adult heart, we kept transgenic mice on a Dox-containing diet until weaning to shut off the expression system, and then switched them to normal lab chow to thereafter induce transgene expression (Fig 3A). Overexpression of zDHHC3 for the first time in the adult heart did not result in immediate bradycardia and dilated cardiomyopathy (data not shown) as observed with perinatal overexpression of zDHHC3 in cardiomyocytes (Fig 2, Supplemental Fig 2). However, zDHHC3 overexpression in adult cardiomyocytes resulted in lethality within 7 or 10 months of transgene expression in the high and low-expressing lines, respectively (Fig 3B). Mortality in DTgZdhhc3 mice was preceded by clinical symptoms of congestive heart failure, including dyspnea and peripheral edema (Fig 3C) as well as cardiac hypertrophy and ventricular and atrial dilation (Fig 3C, D). Cardiac functional assessment by echocardiography revealed left ventricular dilation (Fig 3E), systolic impairment (Fig 3F), and cardiac dysfunction (Fig 3G) in DTgZdhhc3 mice prior to mortality. Lower-expressing lines exhibited an identical phenotype with a delayed onset and progression of disease (Fig 3C-G, Supplemental Table 2). In contrast, overexpression of the zDHHC3^DHHS^ transferase-dead mutant in the adult heart did not cause cardiac hypertrophy, adverse remodeling, or cardiomyopathy at any age examined (Figs 3D-G, Supplemental Table 2). Taken together, these data demonstrate induction of zDHHC3 S-acyltransferase activity in adult cardiomyocytes causes congestive heart failure.

**Figure 3.**
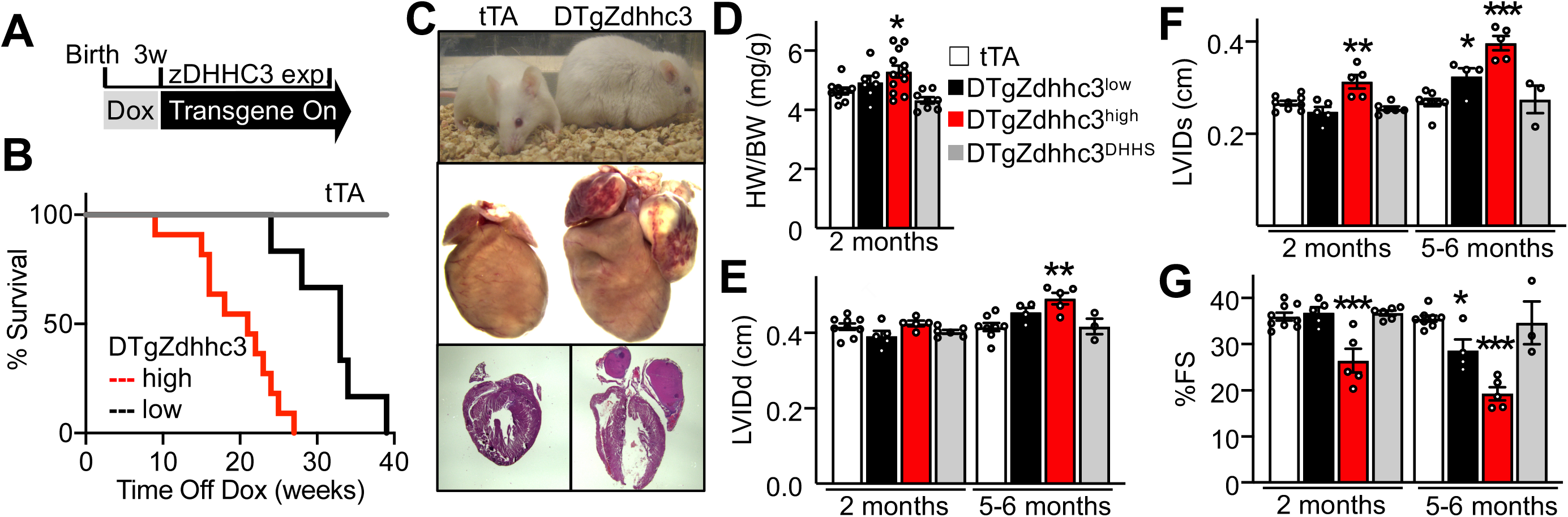
Overexpression of zDHHC3 in the adult heart results in congestive heart failure (CHF). (A) Schematic of transgene expression. Mice were kept on Dox food until weaning and then switched to lab chow to induce transgene expression. (B) Kaplan-Meier survival curve. n= 9 tTA, 6 DTgZdhhc3 ^low^, 11 DTgZdhhc3 ^high^. (C) DTgZdhhc3 mouse and littermate tTA control (low line), whole hearts, and H&E-stained cardiac sections after 7 months of transgene expression. DTg mice exhibit clinical signs of CHF. (D) Heart weight-to-body-weight ratios after 2 months of transgene expression. n=7-12. (E-G) Echocardiographic measurement of (E) diastolic left ventricular inner diameter in diastole (LVIDd), (F) systolic LVID (LVIDs), and (G) % fractional shortening (FS) at the indicated time after transgene expression. The high line of TgZdhhc3 mice were assayed by echocardiography at 5 months after transgene expression due to excessive mortality by 6 months as shown in B. n=3-9. *P<0.05, **P<0.005, ***P<0.0001 compared to tTA, ANOVA with Dunnett’s multiple comparisons test.

To identify zDHHC3 substrates that could underlie cardiac maladaptation in response to enhanced zDHHC3 activity (Figs 1-3), we employed a quantitative and site-specific proteomic approach to sequence peptides containing S-palmitoylated cysteine residues (Fig 4A). We generated stable 3T3 cell lines that overexpress zDHHC3 or GFP as a control and performed stable isotope labeling with amino acids in cell culture (SILAC) for quantitative mass spectrometry sequencing (Fig 4A) ^69^. S-palmitoylated proteins were purified from 3T3-zDHHC3 and 3T3-GFP cells by Acyl Resin-Assisted Capture (Acyl-RAC) ^43^, trypsin-digested on thiopropyl sepharose, and eluted to release bound peptides containing the S-palmitoylation sites for mass spectrometry sequencing (Fig 4A). 3T3-zDHHC3 cells were labeled with “heavy” lysine and arginine while 3T3-GFP controls were labeled with media containing normal isotopic lysine and arginine (“light”) such that peptides identified with increased heavy:light ratios (H:L) exhibit increased S-palmitoylation in zDHHC3 overexpressing cells. We identified 82 unique proteins and 101 unique peptides containing H:L ratios above 1.2 (Fig 4B, data not shown), suggesting regulation by zDHHC3. Peptides sequenced included the previously reported zDHHC3 modification sites on phosphatidylinositol 4-kinase IIα (PI4K2α) ^70, 71^ as well as previously reported S-palmitoylation sites on caveolin-2 ^72^, Rac1 ^17^, scribble ^73^, and Trappc3 ^74^ (Fig 4B, data not shown).

**Figure 4.**
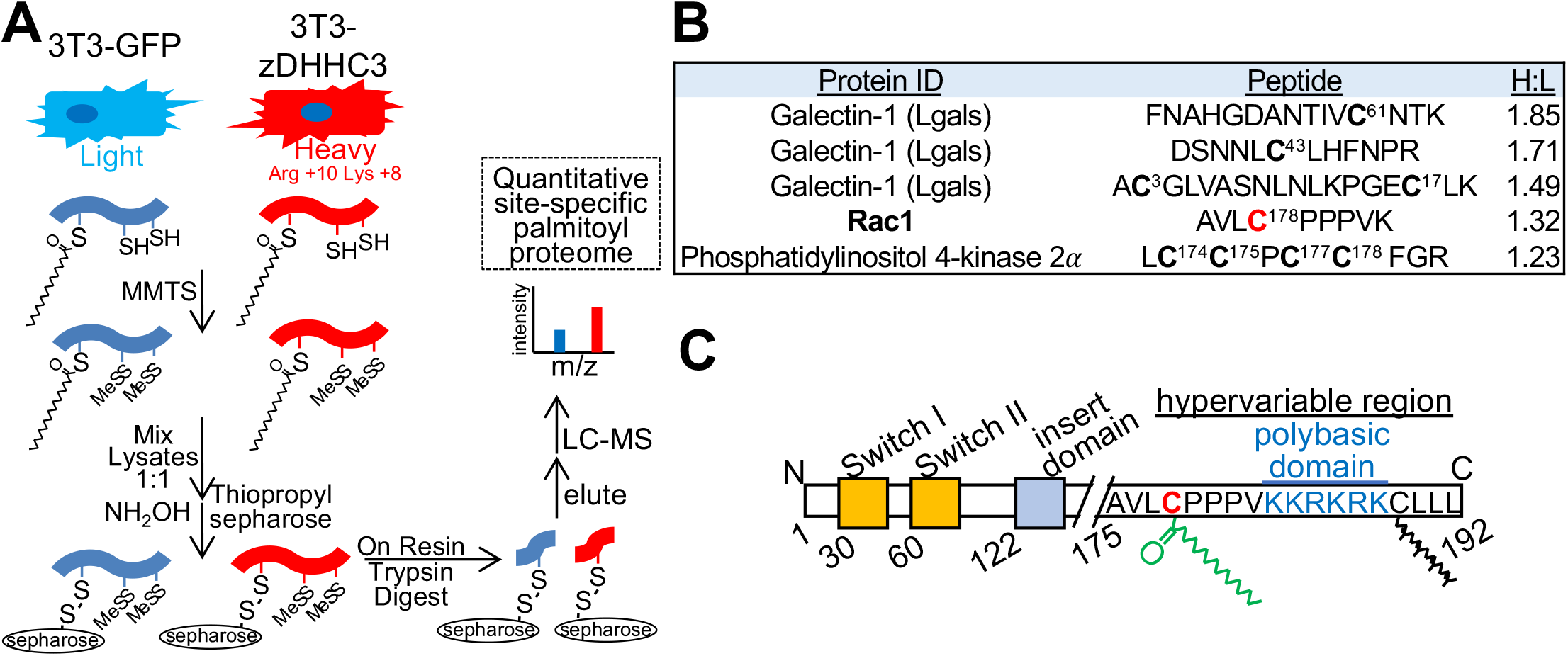
Identification of Rac1 as a novel substrate of zDHHC3. (A) Experimental design for quantitative site-specific S-acyl proteomic identification of zDHHC3 substrates. Stable 3T3 cell lines overexpressing zDHHC3 or GFP as a control were labeled by SILAC and equal protein from “heavy” and “light” cells was mixed and subjected to Acyl Resin-Assisted Capture (Acyl-RAC). S-palmitoylated proteins bound to thiopropyl sepharose were trypsin digested and bound peptides containing S-palmitoylation sites eluted for mass spectrometry sequencing. Adapted from Ref [43]. (B) Table of selected peptides and proteins enriched in 3T3-zDHHC3 cells compared to 3T3-GFP controls with known functions in cardiac hypertrophy or established zDHHC3 substrates. Peptide sequenced from Ras-related C3 botulinum toxin substrate 1 (Rac1) was amongst the top putative zDHHC3-modified peptides with heavy:light ratio (H:L) >1.0. For peptides sequenced multiple times the mean H:L ratio is shown. (D) Diagram of Rac1 protein domain structure including C-terminal hypervariable region that contains membrane targeting motifs, including Cys-178 S-palmitoylation site *(red* Cys, *green* lipid modification), polybasic domain (*blue*), and C-terminal Cys-189 geranylgeranylation (prenylation) site *(black* lipid modification). MMTS, methyl methanethiosulfonate.

Our proteomic screen suggests zDHHC3 directly modifies Rac1 at Cys-178 (Fig 4B), which is critical for its activation and localization to specific plasma membrane microdomains involved in actin cytoskeletal reorganization ^17^. Cys-178 of Rac1 is located in its C-terminal membrane-targeting domain that also contains the classical prenylated -CAAX motif required for processing and trafficking of all small GTPases (Fig 4C) ^75^. Thus, Rac1 undergoes reversible S-palmitoylation at Cys-178 that can regulate membrane microdomain targeting in addition to its permanent generanylgeranyl modification at Cys-189 (Fig 4C) thereby enabling dynamic regulation and diversification of Rac1-mediated signaling. Importantly, S-palmitoylation-dependent regulation of Rac1 has not been evaluated in cardiomyocytes or in vivo to date. To determine if zDHHC3 S-palmitoylates Rac1 in the heart, we performed Acyl-RAC assays to purify S-palmitoylated proteins from transgenic hearts followed by immunoblotting, where we observed a substantial increase in S-palmitoylated Rac1 in zDHHC3 overexpressing hearts (Figs 5A-C, Supplemental Fig 3C, D) concomitant with upregulation of total Rac1 protein levels (Fig 5B, D, Supplemental Fig 3C). H-Ras S-palmitoylation was reduced in zDHHC3 overexpressing hearts (Fig 5B), indicating specificity of zDHHC3 for modification of Rac1 in cardiomyocytes. Notably, induction of Rac1 S-palmitoylation in DTgZdhhc3 hearts occurred within 2 weeks of transgene expression, prior to the development of cardiac hypertrophy and heart failure (Supplemental Fig 3A-D), suggesting modification of Rac1 is a proximal mechanism underlying zDHHC3 activity-induced cardiac pathology. zDHHC3-mediated S-palmitoylation of Rac1 elicited a dramatic enhancement of Rac1 sarcolemmal localization (Fig 5E, F). Immunoblotting of membrane fractions from transgenic hearts demonstrated a substantial increase in membrane-associated Rac1 with zDHHC3 overexpression but not with the enzymatically-dead mutant (Fig 5E). Finally, immunostaining of isolated myocytes from transgenic hearts similarly revealed a dramatic enhancement in plasma membrane-associated Rac1 in zDHHC3-overexpressing cardiomyocytes relative to tTA and DTgZdhhc3 ^DHHS^ controls (Fig 5F).

**Figure 5.**
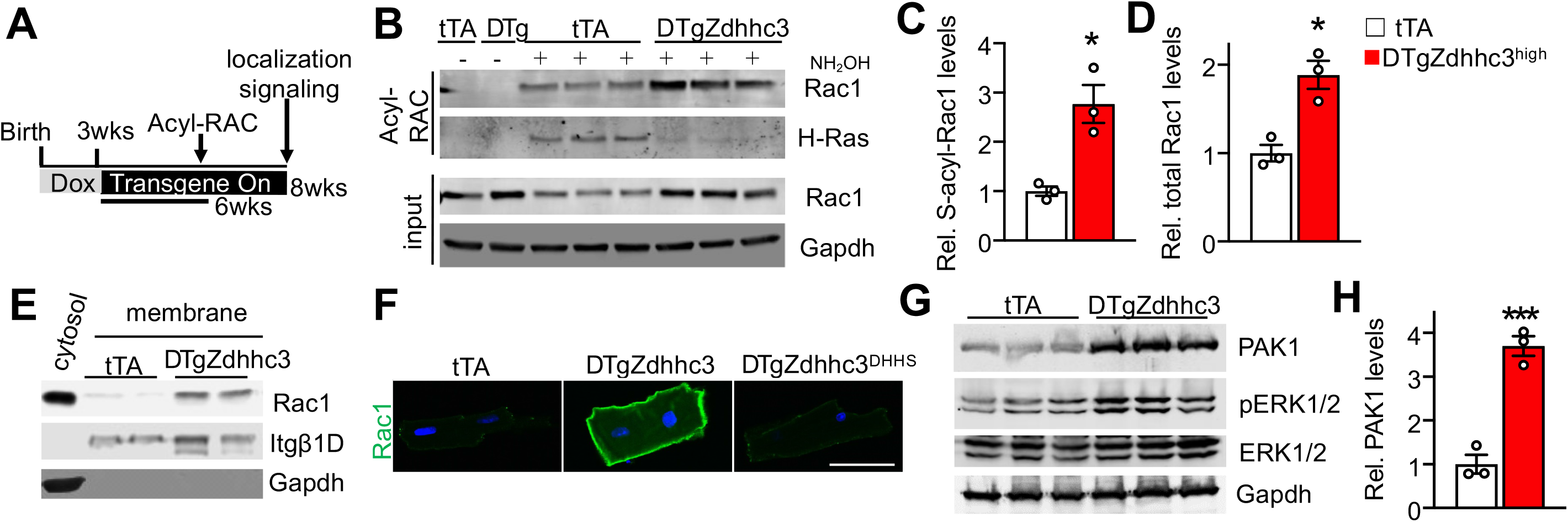
Rac1 S-palmitoylation, membrane localization, and effector signaling are regulated by zDHHC3 in the heart. (A) Experimental design. Hearts were harvested after 6 weeks of transgene expression in the adult heart for analyses of protein S-palmitoylation and after 8 weeks for membrane preparations, immunolocalization, and signaling studies. (B) Immunoblotting for the indicated S-palmitoylated proteins purified by Acyl-RAC. (-) indicates negative controls lacking NH_2_OH treatment. (C-D) Quantification of (C) S-palmitoylated and (D) total Rac1 protein levels normalized to Gapdh from B. n=3. (E) Immunoblotting for the indicated membrane proteins isolated from transgenic hearts. Itgβ1D, integrin β1D. (F) Immunocytochemistry for endogenous Rac1 in cardiomyocytes isolated from transgenic hearts expressing zDHHC3 or the enzymatically-dead zDHHC3^DHHS^ mutant. Scale bar = 50 **μ**m. (G) Immunoblotting for the Rac1 effector PAK1 and phosphorylation of its substrate, ERK1/2, in transgenic hearts and (H) quantification of PAK1 protein levels. n=3. *P<0.05, ***P=0.001 compared to tTA, unpaired t-test.

Western blotting analyses of signaling molecules downstream of Rac1 revealed a substantial increase in the expression of the Rac1 effector, p21-activated kinase 1 (PAK1) ^76, 77^, in zDHHC3 overexpressing hearts (Fig 5G, H) as well as increased phosphorylation of extracellular signal-regulated kinases 1 and 2 (ERK1/2) (Fig 5G), which are activated by PAK1 ^78-80^ and function as transducers of cardiac hypertrophy ^81-83^. Thus, zDHHC3-mediated S-palmitoylation enhances Rac1 translocation to the sarcolemma and downstream activation of PAK1 and ERK1/2.

To further probe the impact of zDHHC3 on Rac1 signaling activity in the heart, we evaluated Rac1 activation in transgenic hearts. DTgZdhhc3 hearts exhibited a substantial increase in the levels of active, GTP-bound Rac1, as well as elevated total levels of Rac1 protein (Fig 6A), indicating zDHHC3-mediated S-palmitoylation of Rac1 (Fig 5B, C, Supplemental Fig 3) induces Rac1 activation and stability. These data are consistent with enhanced Rac1 membrane translocation (Fig 5E, F) and effector signaling (Fig 5G, H) also observed in zDHHC3 overexpressing hearts. Surprisingly, zDHHC3 overexpression had a profound effect on all RhoGTPase family proteins, eliciting an increase in the abundance of not just Rac1 but also RhoA, Cdc42, and RhoGDI (Fig 6A, C). There was also a concomitant elevation in levels of active, GTP-bound RhoA in addition to Rac1 (Fig 6A). RasGTPase expression was unaffected by zDHHC3 activity (Fig 6A, B) suggesting specificity for RhoGTPase signaling. Indeed, protein levels of Rac1, RhoA, and Cdc42, but not H-Ras, were elevated in both cytoplasmic and membrane fractions isolated from zDHHC3 overexpressing hearts compared to tTA controls (Fig 6B). RhoGDI serves as a master regulator of RhoGTPase signaling homeostasis by regulating the abundance, activity, and localization of all RhoGTPase family proteins ^11, 12, 84^. Here we observed that protein levels of RhoGDI were also substantially increased in zDHHC3 overexpressing hearts (Fig 6A, C), suggesting a broad-spectrum effect of zDHHC3 activity on RhoGTPase signaling. This remarkable induction of RhoGTPases and RhoGDI did not occur in hearts overexpressing the zDHHC3^DHHS^ transferase-dead mutant (Fig 6C), indicating zDHHC3 S-acyltransferase activity directly underlies the observed increase in RhoGTPase signaling.

**Figure 6.**
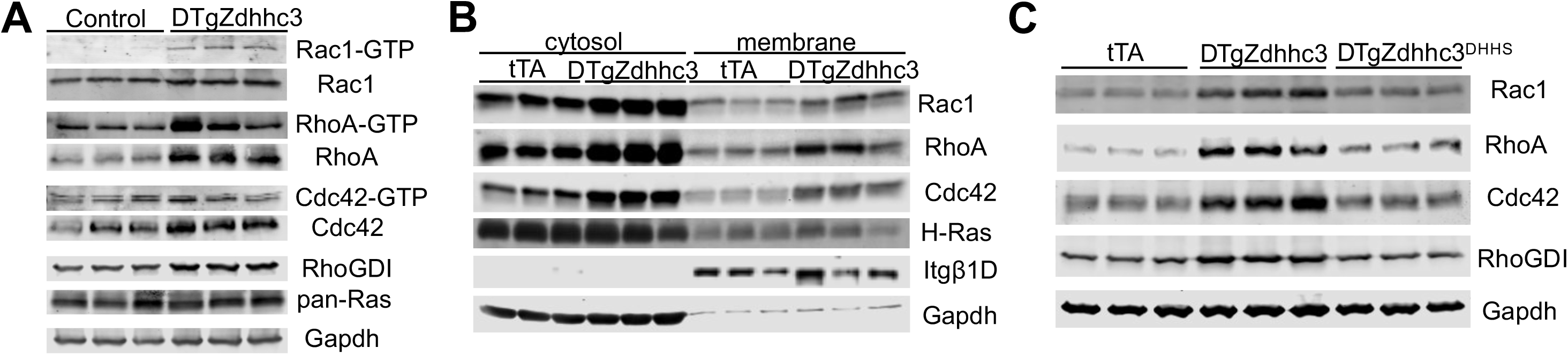
Enhanced zDHHC3 activity induces signaling by all Rho family small GTPases in the heart. Immunoblotting for (A) active and total levels of Rho family small GTPases in transgenic hearts overexpressing zDHHC3 (low line, 5 months of transgene expression in adult heart as in Fig 3), (B) cytosolic and membrane protein fractions isolated from transgenic hearts, and (C) total RhoGTPase protein levels in hearts overexpressing zDHHC3 or the enzymatically-dead zDHHC3^DHHS^ mutant (high line, 2 months of transgene expression in the adult heart). Controls in A are nontransgenic or single transgenic tTA or Zdhhc3 littermates of DTgZdhhc3 mice overexpressing zDHHC3.

## DISCUSSION

S-palmitoylation plays critical roles in the pathophysiology of cancer ^45-49, 85^, inflammation ^50-53^, peripheral artery disease ^86^, and thrombosis ^87^, yet no direct role has been established for this post-translational modification in the pathogenesis of cardiac hypertrophy and heart failure. Indeed, despite S-palmitoylation of essential cardiac signal transducing proteins (i.e. α- and β-adrenergic receptors ^88, 89^, endothelin receptors ^90-92^, Gαq ^93, 94^, and Gαs ^94, 95^), little is known about its functions in the heart beyond its roles in ion channel regulation ^40, 54, 96^. Here, we surveyed zDHHC S-acyltransferase enzymes to uncover novel regulators of cardiomyocyte signal transduction and homeostasis and found that activity of zDHHC3 and zDHHC7 at the cytoplasmic surface of the Golgi promotes hypertrophic signaling and cardiomyopathy in vivo. Further investigation identified Rac1 as a novel substrate of zDHHC3 through an unbiased proteomic screen, and upregulation of Rac1 S-palmitoylation and signaling activity preceded heart failure in zDHHC3 transgenic mice, showing a relationship between zDHHC3-regulated S-palmitoylation of Rac1 and maladaptive hypertrophic signaling.

Rac1 plays fundamental roles in cardiac homeostasis and pathophysiology and is necessary and sufficient to induce cardiac hypertrophy ^5, 21, 97^ and arrhythmia ^98, 99^. Overexpression of constitutively active Rac1 results in lethal dilated cardiomyopathy ^21^ and arrhythmogenesis ^98^ while loss of Rac1 in cardiomyocytes ameliorates angiotensin II-induced cardiac hypertrophy and oxidative stress ^5^. Rac1 canonically signals from lamellipodia and membrane ruffles in non-muscle cells ^77, 100, 101^ but how Rac1 functions in cardiomyocytes with a relatively static cytoskeleton, limited migration and proliferation, and unique sarcolemmal signaling domains is not well understood. Here, we uncovered a novel regulatory mechanism governing cardiomyocyte Rac1 signaling activity through zDHHC3-mediated S-palmitoylation.

Functions of small GTPases are regulated by their translocation to cellular membranes where they modulate signaling by effector molecules that ultimately impact a host of cellular processes including cell growth, proliferation, and migration ^13, 102^. All Ras superfamily small GTPases (which also includes Rho and Rab family GTPases) undergo prenylation on their C-terminus, the irreversible modification of cysteines with an unsaturated isoprenyl fatty acid, which is critical for their processing, trafficking, and ultimately membrane association ^103-105^. Rho family GTPases (RhoA/C, Rac1, Cdc42) are primarily regulated by RhoGDI, a Rho-specific chaperone molecule that binds the prenylated C-terminus of RhoGTPases to regulate their delivery to and extraction from sites of action at cell membranes ^11, 12, 84^. We uncovered zDHHC3-regulated S-palmitoylation of Rac1 at Cys-178, adjacent to its C-terminal polybasic region and prenylated CAAX motif, which has been shown to target Rac1 to higher ordered cholesterol-rich membrane microdomains and increase its activation ^17^. These data suggest zDHHC3-mediated Rac1 S-palmitoylation compartmentalizes Rac1 at distinct sarcolemmal signaling microdomains that likely underlie pathological remodeling and hypertrophy. zDHHC3 overexpressing hearts exhibit remarkable induction of the Rac1 effector kinase, PAK1, and phosphorylation of ERK1/2, regulators of hypertrophic cardiac growth ^78, 80-82^. These data collectively suggest a working hypothesis whereby zDHHC3 activity at the cardiomyocyte Golgi S-palmitoylates Rac1 to promote its sarcolemmal translocation and signaling activity along with induction of all small GTPases of the Rho family and RhoGDI, which is associated with congestive heart failure in zDHHC3 transgenic mice that also phenocopies cardiac-specific overexpression of RhoA or Rac1 ^9, 21^.

S-palmitoylation can function as a dynamic and powerful post-translational regulatory mechanism to rapidly alter intracellular signal transduction ^28, 54, 75^. Soluble signaling proteins, such as Ras family small GTPases and G-protein α subunits, can undergo inducible S-palmitoylation to facilitate translocation to select membrane domains where they associate with receptors and activate signaling effectors followed by depalmitoylation and S-palmitoylation again by Golgi-localized zDHHC enzymes. This results in successive rounds of membrane translocation and sustained signaling output in a process termed palmitoylation cycling ^31, 32, 50, 106, 107^. For example, N-Ras is S-palmitoylated at the Golgi by zDHHC9 and depalmitoylated at the plasma membrane by ABHD17 family thioesterases to dynamically regulate N-Ras signaling activity. Consequently, loss of zDHHC9 or inhibition of ABHD17 enzymes impairs growth and proliferation of N-Ras-dependent cancer cells ^16, 85, 107-109^. Similarly, STAT3 is S-palmitoylated by zDHHC7 at the Golgi to induce its association with growth factor receptors and phosphorylation at the membrane where it is then depalmitoylated by APT-2 to enable nuclear translocation and activation of an inflammatory gene expression program ^50^. Disruption of STAT3 S-palmitoylation cycling by genetic deletion of zDHHC7 or pharmacological inhibition of APT-2 ameliorates inflammatory gene expression and colitis in an animal model inflammatory bowel disease ^50^. Thus, palmitoylation cycling of soluble signaling proteins can provide a regulatory mechanism to garner sustained signaling and activation of downstream transduction circuitry. Here, we observed that zDHHC3 activity in cardiomyocytes regulates S-palmitoylation of Rac1 and consequently its translocation to the sarcolemma, GTP-loading, and activation of downstream effectors, suggesting zDHHC3-mediated regulation of Rac1 palmitoylation cycling induces pathogenic signaling, which could represent a new therapeutic vantage point to treat cardiomyopathy.

In summary, our data are the first to demonstrate a critical role for dynamic S-palmitoylation in the heart as a regulator of pathologic signal transduction that leads to hypertrophy and maladaptive ventricular remodeling. Interestingly, statin drugs commonly prescribed to treat cardiovascular disease repress membrane localization, activation, and abundance of Rac1 in cardiomyocytes ^26, 110, 111^ and similarly reduce cardiac Rac1 activity and oxidative stress in human heart failure ^23^. Indeed, the efficacy of statin drugs in heart failure treatment is thought to be mediated in part through repression of Rac1 via inhibition of prenylation and antagonism of maladaptive Rac1 signaling and oxidative stress ^27, 110, 112-114^. Inhibition of zDHHC3 or Rac1 S-palmitoylation may thus provide an alternative therapeutic approach for cardiac disease treatment by inhibiting maladaptive Rac1 signaling at the sarcolemma.

## ACKNOWLEDGEMENTS

This work was supported by the National Heart, Lung, and Blood Institute grant R00HL136695 to MJB.

