## Supplemental Data for "S-Palmitoylation-Dependent Regulation of Cardiomyocyte Rac1 Signaling Activity and Cardiac Hypertrophy"

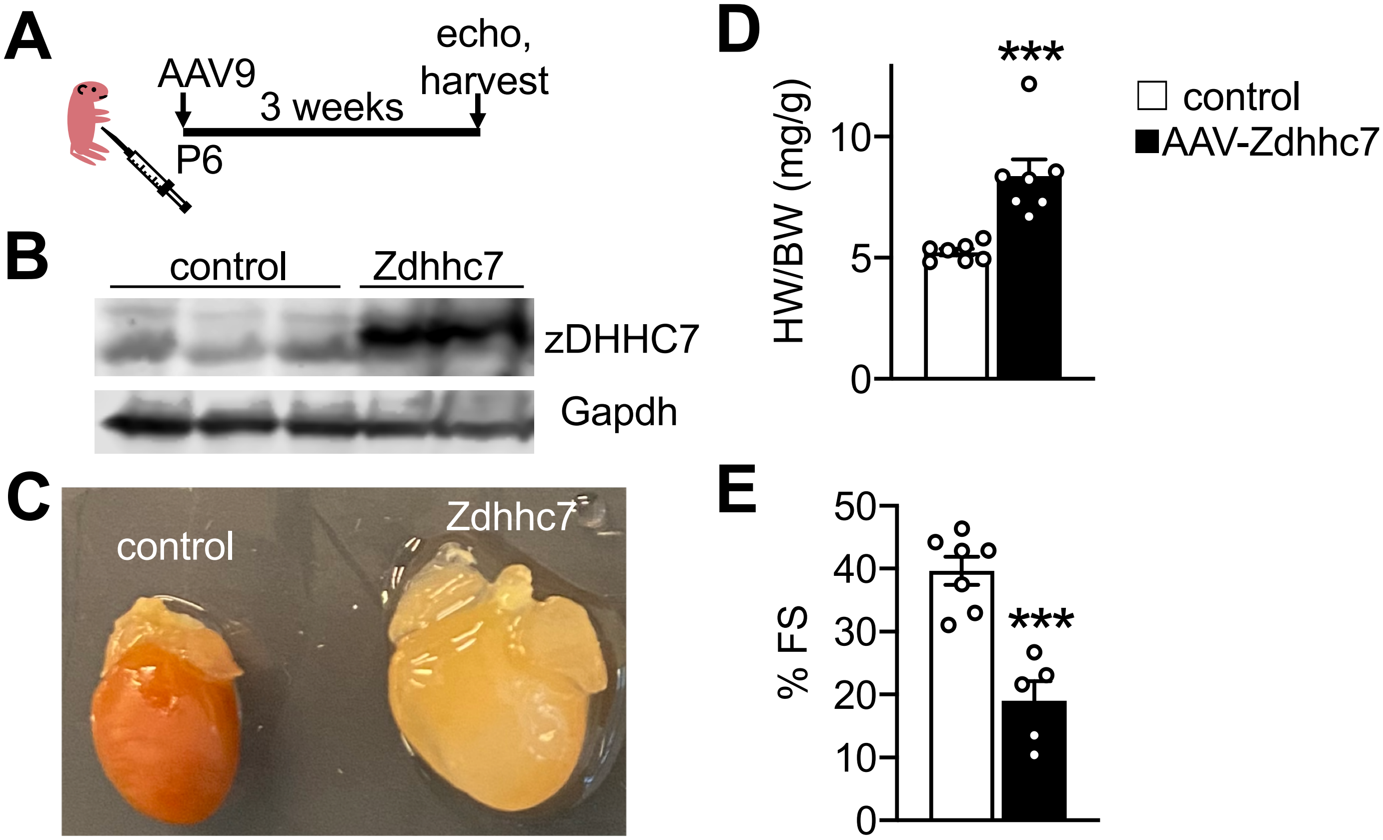


**Supplemental Fig 1**. Overexpression of zDHHC7, the most closely related S-acyl transferase to zDHHC3, also causes cardiomyopathy. (A) Experimental design schematic and (B) Western blotting for AAV9-mediated overexpression of zDHHC7 in the heart. (C) Gross morphology, (D) heart weight-to-body weight ratios, n=7, (E) and fractional shortening (FS) as assessed by echocardiography in mice with cardiac overexpression of zDHHC7, 3 weeks after AAV injection. n=5-7, ***P<0.001, unpaired t-test.

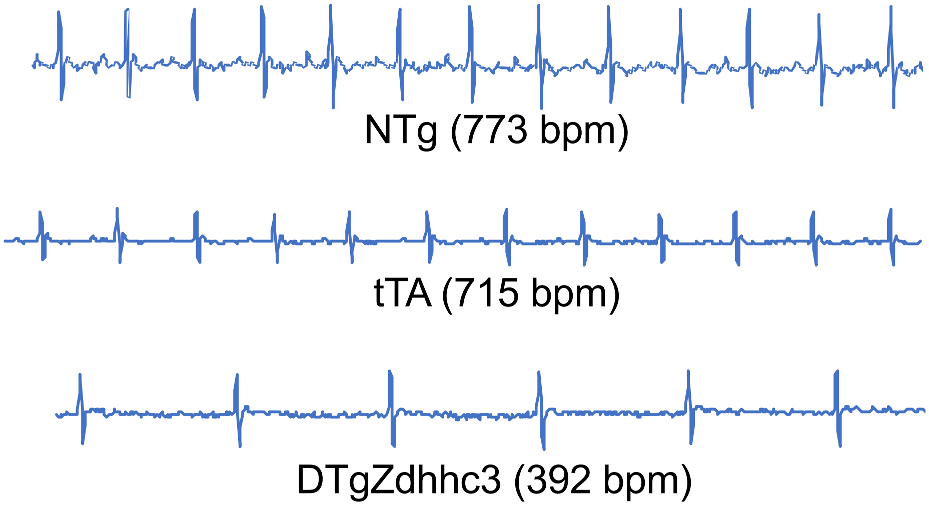


**Supplemental Fig 2**. Severe bradycardia in mice with cardiomyocyte-specific overexpression of zDHHC3 from birth. Electrocardiography was performed on conscious mice from the low-expressing line of TgZdhhc3 mice at 5 weeks of age. Representative electrocardiograms (ECGs) of mice of the indicated genotypes are shown. n= 2 controls and 2 DTg. NTg, nontransgenic. bpm, beats per minute.

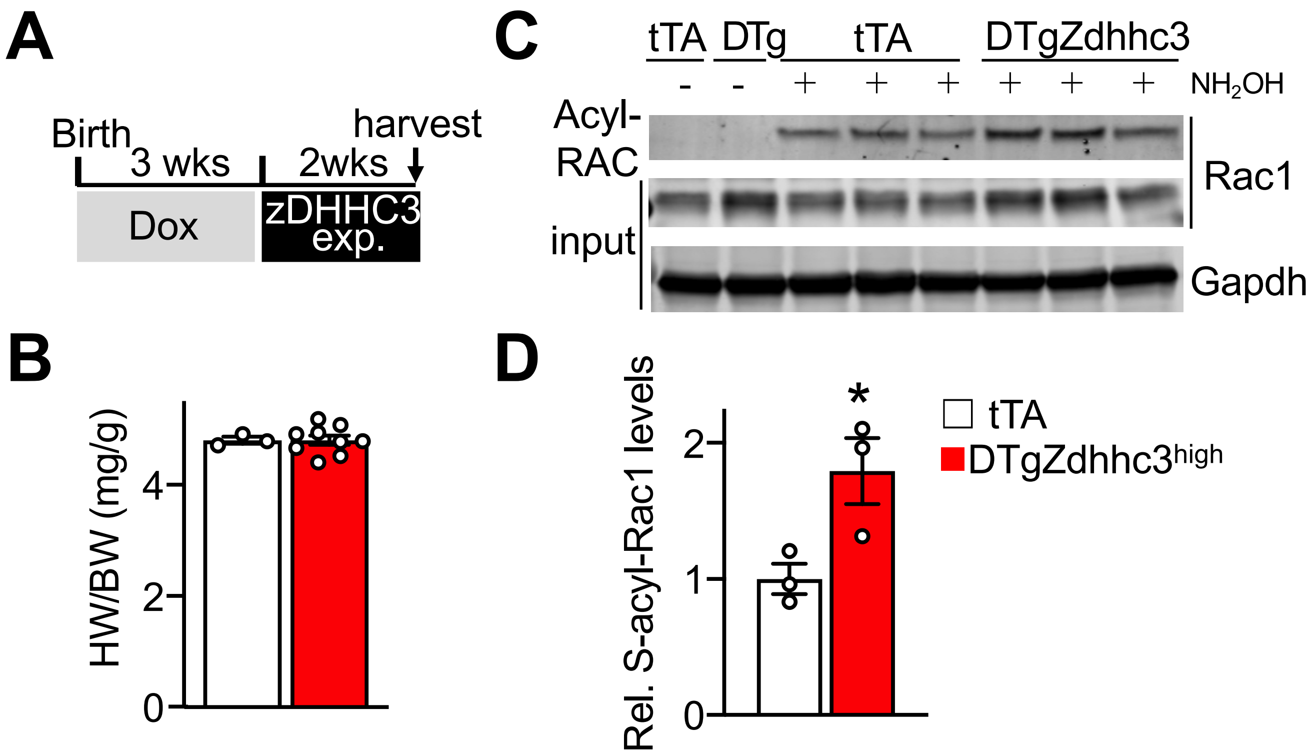


**Supplemental Fig 3**. Induction of Rac1 S-palmitoylation precedes cardiac hypertrophy and failure in TgZdhhc3 mice (high line). (A) Experimental design. Hearts were harvested after 2 weeks of transgene expression in the adult heart for analyses of protein S-palmitoylation. (B) Heart weight-to-body weight ratios. n=3-9. (C) Immunoblotting for S-palmitoylated and total Rac1 in transgenic hearts. (D) Quantification of S-palmitoylated Rac1 normalized to Gapdh expression. n=3. *P<0.05 compared to tTA, unpaired t-test.

| Genotype | tTA | DTgZdhhc3 (low) | DTgZdhhc3$^{DHHS}$ |
|---|---|---|---|
| N | 10 | 4 | 8 |
| IVSd (cm) | 0.068 ± 0.003 | 0.074 ± 0.003 | 0.080 ± 0.005* |
| LVIDd (cm) | 0.402 ± 0.008 | 0.616 ± 0.031*** | 0.407 ± 0.009 |
| LVPWd (cm) | 0.093 ± 0.003 | 0.108 ± 0.007 | 0.096 ± 0.004 |
| IVSs (cm) | 0.117± 0.004 | 0.108 ± 0.008 | 0.125 ± 0.005 |
| LVIDs (cm) | 0.254 ± 0.007 | 0.502 ± 0.045*** | 0.263 ± 0.010 |
| LVPWs (cm) | 0.121± 0.004 | 0.130 ± 0.004 | 0.124 ± 0.004 |
| RR int (sec) | 0.140 ± 0.005 | 0.351 ± 0.141** | 0.157 ± 0.008* |
| %FS | 36.70 ± 0.89 | 18.95 ± 4.05*** | 35.38 ± 1.19 |

**Supplemental Table 1.** Echocardiographic assessment of cardiac structure and function in Zdhhc3 transgenic mice and transgenic mice expressing the enzymatically dead zDHHC3$^{DHHS}$ mutant at 6-8 weeks of age. **α**-myosin heavy chain promoter-driven Zdhhc3 transgene expression begins around birth in ventricular myocytes as Figs 2D-I. tTA, tetracycline transactivator; DTg, double transgenic; d, diastolic; s, systolic; IVS, interventricular septal thickness; LVID, left ventricular (LV) inner diameter; LVPW, LV posterior wall thickness; FS, fractional shortening; int, interval. *P<0.05, **P<0.01, ***P<0.0001 compared to tTA controls at the same timepoint, ANOVA with Dunnett's multiple comparisons test.

| Time off Dox | 2 months | | | | 5-6 months | | | |
|---|---|---|---|---|---|---|---|---|
| Genotype | tTA | DTgZdhhc3 (low) | DTgZdhhc3 (high) | DTgZdhhc3$^{DHHS}$ (high) | tTA | DTgZdhhc3 (low) | DTgZdhhc3 (high) | DTgZdhhc3$^{DHHS}$ (high) |
| N | 9 | 5 | 5 | 6 | 8 | 4 | 5 | 3 |
| IVSd (cm) | 0.083 ± 0.002 | 0.076 ± 0.004 | 0.077 ± 0.002 | 0.078 ± 0.003 | 0.082 ± 0.004 | 0.079 ± 0.006 | 0.086 ± 0.007 | 0.093 ± 0.004 |
| LVIDd (cm) | 0.416 ± 0.008 | 0.392 ± 0.014 | 0.425 ± 0.007 | 0.403 ± 0.005 | 0.415 ± 0.011 | 0.454 ± 0.013 | 0.491 ± 0.015** | 0.417 ± 0.021 |
| LVPWd (cm) | 0.104 ± 0.003 | 0.099 ± 0.004 | 0.096 ± 0.004 | 0.102 ± 0.004 | 0.103 ± 0.003 | 0.108 ± 0.002 | 0.106 ± 0.007 | 0.099 ± 0.007 |
| IVSs (cm) | 0.128 ± 0.004 | 0.122 ± 0.004 | 0.111 ± 0.004* | 0.135 ± 0.004 | 0.126 ± 0.002 | 0.120 ± 0.003 | 0.114 ± 0.005 | 0.133 ± 0.010 |
| LVIDs (cm) | 0.266 ± 0.005 | 0.248 ± 0.011 | 0.313 ± 0.015** | 0.255 ± 0.005 | 0.268 ± 0.008 | 0.325 ± 0.018* | 0.397 ± 0.015*** | 0.274 ± 0.030 |
| LVPWs (cm) | 0.133 ± 0.004 | 0.125 ± 0.006 | 0.127 ± 0.004 | 0.124 ± 0.004 | 0.143 ± 0.003 | 0.135 ± 0.006 | 0.122 ± 0.006** | 0.129 ± 0.004 |
| RR int (sec) | 0.138 ± 0.00 | 0.147 ± 0.006 | 0.143 ± 0.008 | 0.163 ± 0.017 | 0.113 ± 0.003 | 0.144 ± 0.010** | 0.143 ± 0.007** | 0.128 ± 0.007 |
| %FS | 35.97 ± 0.85 | 36.84 ± 1.19 | 26.44 ± 2.57*** | 36.75 ± 0.52 | 35.55 ± 0.67 | 28.60 ± 2.45* | 19.26 ± 1.43*** | 34.63 ± 4.64 |

**Supplemental Table 2.** Echocardiographic assessment of cardiac structure and function in lines of Zdhhc3 transgenic mice. Transgenic mice on doxycycline chow (Dox) until weaning were placed on normal lab chow to induce transgene expression in the adult heart for the indicated time as described in Fig 3. tTA, tetracycline transactivator; DTg, double transgenic; d, diastolic; s, systolic; IVS, interventricular septal thickness; LVID, left ventricular (LV) inner diameter; LVPW, LV posterior wall thickness; FS, fractional shortening; int, interval. *P<0.05, **P<0.01, ***P<0.0001 compared to tTA controls at the same timepoint, ANOVA with Dunnett's multiple comparisons test..
